# Differentiable Search of Evolutionary Trees

**DOI:** 10.1101/2023.07.23.550206

**Authors:** Ramith Hettiarachchi, Avi Swartz, Sergey Ovchinnikov

## Abstract

Inferring the most probable evolutionary tree given leaf nodes is an important problem in computational biology that reveals the evolutionary relationships between species. Due to the exponential growth of possible tree topologies, finding the best tree in polynomial time becomes computationally infeasible. In this work, we propose a novel differentiable approach as an alternative to traditional heuristic-based combinatorial tree search methods in phylogeny. The optimization objective of interest in this work is to find the most parsimonious tree (i.e., to minimize the total number of evolutionary changes in the tree). We empirically evaluate our method using randomly generated trees of up to 128 leaves, with each node represented by a 256-length protein sequence. Our method exhibits promising convergence (< 1% error for trees up to 32 leaves, < 8% error up to 128 leaves, given only leaf node information), illustrating its potential in much broader phylogenetic inference problems and possible integration with end-to-end differentiable models. The code to reproduce the experiments in this paper can be found at https://github.ramith.io/diff-evol-tree-search.

## 1 Introduction

Evolutionary trees (or phylogenetic trees) provide biologists with a structured, hierarchical representation of how current species are related through hypothetical ancestors that are probably extinct at present. Beyond theoretical constructs, they have practical applications in various fields of biology and medicine. For instance, phylogenetic techniques are crucial in the decision-making process when responding to emerging viruses Attwood et al. (2022).

Parsimony methods are one of many methods (such as distance methods, maximum likelihood, maximum compatibility) for constructing evolutionary trees. The parsimony principle states that the most acceptable explanation of an occurrence is the one that requires the minimum number of assumptions or explanations Sober (1981). Thus, inferring the most parsimonious tree given leaf nodes requires finding the tree topology and its corresponding ancestral representations that explains the data with minimum number of evolutionary steps. This combinatorial problem was shown to be NP-Complete Foulds & Graham (1982); Steel (1992).

Due to the complexity of the problem, existing methods consider heuristic search techniques by limiting the search space. Although this does not guarantee that the algorithm will find the optimal solution, they do facilitate exploration of a vast number of tree topologies, starting from an initial guess and iteratively refining it. These methods can be broadly categorized into 1) tree rearrangement methods 2) branch and bound methods 3) neighbor joining methods Saitou & Nei (1987); Giribet (2007); Felsenstein (2004).

With the growth of deep learning methods, there have been several new directions in constructing evolutionary trees. Zhu & Cai (2021) et al. propose an alignment-free method in which an attention model Vaswani et al. (2017) is trained through reinforcement learning to reconstruct evolutionary trees. However, this method requires algorithmic post-processing to produce the final tree, thus, prevents it from being end-to-end differentiable. Azouri et al. (2023) demonstrate a deep-Q-learning agent on empirical data consisting of up to 20 leaves.

With the success of using hyperbolic geometry for hierarchical data, there has been work on obtaining continuous embeddings for trees Monath et al. (2019); Chami et al. (2020); Corso et al. (2021). Subsequently, optimization in the hyperbolic space for phylogeny Wilson (2021), developing new metrics Matsumoto et al. (2021) and addressing the phylogenetic placement problem Jiang et al. (2022) have been explored. Given the challenge of scalability in traditional Bayesian phylogenetics, methods based on variational inference have been proposed Zhang & Matsen (2019); Dang & Kishino (2019). Recently, Zhang (2023) proposed a topological feature learning framework for phylogenetic inference using graph neural networks.

In contrast to previous work, our approach circumvents the discreteness of the raw tree and sequence representations in the first place and models their relationship in a differentiable manner. By doing so, we obtain a soft-parsimony score that can be optimized in an end-to-end differentiable manner, without the need for any prior training data.

We perform experiments for tree topologies up to 128 leaves and analyze our method for 3 tasks. 1) find the tree given all sequences, 2) find the ancestors given the tree topology and leaves (small parsimony), and 3) find the tree and the ancestors given leaves (maximum parsimony). For the small parsimony problem we achieve the ≈0% mean error for all leaf counts experimented, meaning that our approach can find the optimal ancestral sequences if the tree topology is known. For the maximum parsimony problem, we achieve < 1% error for trees up to 32 leaves, < 8% error up to 128 leaves.

Our work opens up new realms for integration with models with more complex cost functions that go beyond site-independence assumption. For example, the cost function can integrate pseudo-likelihood between nodes using protein language models or those conditioned on protein structure.

## 2 Notation and background

### 2.1 Maximum parsimony problem (find tree and ancestors)

A rooted phylogenetic tree is a directed acyclic graph (DAG) *G* = (*V, E*). Given a set of *N* leaves, the maximum parsimony problem intends to find the phylogenetic tree (𝒯) and ancestor nodes that describe the given data with minimum number of evolutionary steps Carmel et al. (2014); Kannan & Wheeler (2012). For a fixed alphabet 𝒜 = {1, …, *c*}, each node in this DAG can be represented by an *l* dimensional vector **s** = (*s*_1_, …, *s*_*l*_) such that **s** ∈ 𝒜^*l*^. We consider the Hamming distance *d* on the node representation *d*((*s*_1_, …, *s*_*l*_), 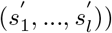 that describes the number of indices *i* such that 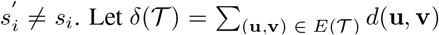 *d*(**u, v**) represent the total number of evolutionary changes in the tree 𝒯. The maximum parsimony problem is then to find the tree that minimizes *δ*(𝒯).

### 2.2 Complexity

The maximum parsimony problem has been comprehensivey studied in the literature as a special case of the Steiner tree problem Hwang & Richards (1992). Further Foulds & Graham (1982) showed that even when |𝒜| = 2, the problem of finding the tree with the minimum number of evolutionary steps is NP-Complete. Given *N* leaves, the number of rooted bifurcating tree topologies that exists can be calculated as (2*N* −3)!! Cavalli-Sforza & Edwards (1967). Thus, even for a tree with only 12 leaves, there are more than 13 billion tree topologies in total.

### 2.2 Small parsimony problem (find ancestors)

The small parsimony problem Carmel et al. (2014) is a much simplified version of the problem in which the phylogenetic tree topology 𝒯 is already given. Therefore, the task is to find the best possible ancestors. There are a number of dynamic programming (DP) algorithms proposed to solve this problem in polynomial such as the Fitch’s algorithm Fitch (1971) and the Sankoff’s algorithm Sankoff (1975).

## 3 Methodology

In the following subsections, we consider how the discrete aspects of the problem are relaxed and how gradient-based optimization can be performed.

### 3.1 Relaxations

There are two aspects to this problem that make it inherently non-differentiable. First, each element of the sequence is combinatorial. Second, considering the adjacency matrix, the space of valid and meaningful tree topologies is sparse.

#### Sequence representation

In both the small and maximum parsimony problems, the ancestor sequences are unknown. We denote learnable parameters *ϕ*_*seq*_ ∈ ℝ^(2*N*−1)×*l*×*c*^ to represent the ancestors to be optimized. To obtain a continuous relaxation of the categorical nature of amino acid types, we transform the real tensor into a tensor 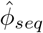, where each element represents a probability distribution over the character space for each position of the sequence. This transformation is done using the softmax function Jang et al. (2017), with the sharpness of the distribution controlled by the temperature parameter *τ*_2_.

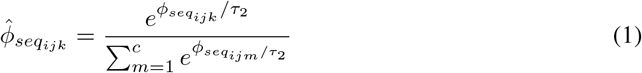

This probability tensor 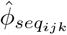 representing ancestors, combined with the one-hot encoding of known leaves results in the node representation tensor *S*.

#### Tree representation

Since the adjacency matrix of any DAG can be permuted to be a strictly upper triangular matrix Nicholson (1975); Li et al. (2022); Charpentier et al. (2022), we ensure the acyclicity of the graphs represented by the adjacency matrix (*A*) by enforcing it to be strictly upper triangular. Furthermore, since leaves cannot be connected to each other, we ensure that the first *N* columns of *A* are zero. The remaining positions are parameterized as *θ*_*T*_.

Due to the irregular structure of the adjacency matrix representation and to speed up the implementation, we first set the non-parameterized region of the adjacency matrix to −inf, and then apply the softmax function for each row. This ensures that the parameterized positions of the adjacency matrix will be represented by the correct probability. Thus, the *i*^*th*^ row of *A* represents the probabilities of node *i* being a child of all other nodes, respectively.

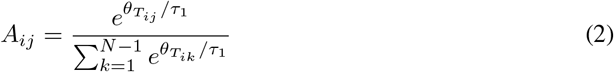

### 3.2 Tree enforcing loss function

With the relaxation of the adjacency matrix, optimizing parameters to reduce *L*_*cost*_ does not explicitly guide the optimization towards a bifurcating tree. Therefore, we enforce the following regularization constraint to maintain the bifurcating property. This regularization forces tree nodes to have exactly two child node connections.

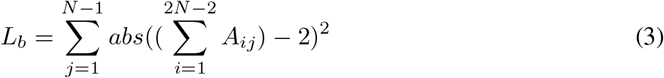

### 3.3 Differentiable soft parsimony score calculation

To calculate the number of evolutionary steps that have occurred in the DAG, we formulate the evolutionary cost calculation as follows. The transposed node representation tensor 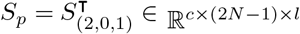 is constructed by representing the characters of the alphabet as its first dimension.

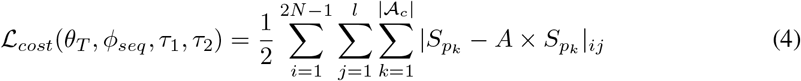

In this equation, the adjacency matrix *A* describes the connection of each node to its parent in the tree structure. The matrix *S*_*pk*_, considers *k*^*th*^ character of the alphabet at a time. Thus, in the matrix *S*_*pk*_, (*S*_*pk*_)_*ij*_ = 1 iff *i*^*th*^ sequence has *j*^*th*^ position equal to the *k*^*th*^ character in the alphabet 𝒜. The matrix multiplication *A* × *S*_*pk*_ serves as a lookup of the sequence table (*S*_*pk*_) and returns the parent corresponding to each node in the tree. Thus, the difference between the matrices |*S*_*pk*_ − *A* × *S*_*pk*_| represents the distance between each child and its parent in the *k*^*th*^ character space. By summing these differences over all dimensions, the cost function captures the overall evolutionary cost. For a visual depiction of this calculation, refer to the Appendix Figure 6.

### 3.4 Bi-level optimization

In the maximum parsimony problem, we need to traverse both the tree and sequence spaces, and there is a dependency between these two. Note that for each tree topology, there is a best set of sequences that define how good each topology is. And once the topology changes, these sequences are no longer valid. Thus, for this task, we formulate the optimization procedure as a bi-level optimization problem. Experimentally, we obtain better results with this formulation than optimizing both of the parameters independently (see Section 4.3 for the ablation study).

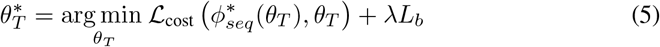

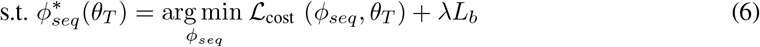

During implementation, we perform *k* gradient descent iterations for the inner objective (Eq. 6). Gradient descent is performed for both objectives using the Adam optimizer Kingma & Ba (2015). Furthermore, we use the JAXopt library to optimize multiple initialization seeds as a batch and obtain the best result Blondel et al. (2021).

### 3.5 Evaluation

In order to evaluate our method, we first generate known evolutionary trees (each consisting of 4− 128 leaves, 256 sequence length, alphabet size *c* = 20). Therefore, we generate complete binary trees starting with a random sequence as the root, make two copies of the sequence at each node and generate two random sets of indices, each with *m* = 50 elements that are mutated to a different character. For each leaf count, we perform 10 random initializations for the leaf sequences to generate examples. Further details on ground truth generation are included in Appendix A.

## 4 Results

We first analyze a simpler task in which all nodes *S* (leaves and ancestors) are known, but the tree topology is unknown. For all leaf counts (*N* = 4 to 128), the adjacency matrix converged to the groundtruth tree. This task is similar to a hierarchical clustering task, where all the nodes are known, and we need to establish a hierarchical dependency between them.

### 4.1 Small Parsimony (find ancestors given tree)

The ground-truth ancestor solutions for the known tree topology is obtained by the Sankoff algorithm. Table 1 shows the results as number of leaves increases. For all cases, the error between ours and the optimal parsimony score is ≈ 0.

**Table 1:** Evaluation on the Small Parsimony Problem.

| Task |  | Small Parsimony (find seqs given tree) |  |  |
| --- | --- | --- | --- | --- |
| $N$ | Mean optimal solution | Mean solution | Mean error | Mean error as a % w.r.t optimal solution |
| 4 | 277.2 | 277.2 | $0.0 \pm 0.0$ | 0.000% |
| 8 | 653.1 | 653.1 | $0.0 \pm 0.0$ | 0.000% |
| 16 | 1407.6 | 1407.6 | $0.0 \pm 0.0$ | 0.000% |
| 32 | 2915.4 | 2915.4 | $0.0 \pm 0.0$ | 0.000% |
| 64 | 5929.3 | 5929.3 | $0.0 \pm 0.0$ | 0.000% |
| 128 | 11971.1 | 11971.3 | $0.2 \pm 0.4$ | 0.001% |

### 4.2 Maximum Parsimony (find ancestors and tree)

In this task, only the leaf sequences are known, and we need to optimize towards the best tree topology and ancestors. As shown in Figure 3 and Table 2, the mean error increases as the number of leaves increases. Note that for up to *N* = 8 the possible tree topologies are even enumerable, as they result in only 135,135 combinations. Thus, our method also converges to optimal solutions. However, from *N* = 16 to 128 leaves, the number of possible tree topologies grows from ≈10^15^ to 10^250^ possibilities and our method converges to local optima. We intend to explore methods to simplify the loss landscape and gradually increase the complexity in order to discover better solutions. Figure 4 depicts the start and end of the optimization process for the maximum parsimony task given randomly initialized *N* = 32 leaves.

**Table 2:** Evaluation on the Maximum Parsimony Problem.

| Task |  | Maximum Parsimony (find both tree & seqs) |  |  |
| --- | --- | --- | --- | --- |
| $N$ | Mean optimal solution | Mean solution | Mean error | Mean error as a % w.r.t optimal solution |
| 4 | 277.2 | 277.2 | $0.0 \pm 0.0$ | 0.000% |
| 8 | 653.1 | 653.1 | $0.0 \pm 0.0$ | 0.000% |
| 16 | 1407.6 | 1407.7 | $0.1 \pm 0.3$ | 0.007% |
| 32 | 2915.4 | 2936.3 | $20.9 \pm 7.4$ | 0.717% |
| 64 | 5929.3 | 6188.6 | $259.3 \pm 27.4$ | 4.373% |
| 128 | 11971.1 | 12885.5 | $914.4 \pm 99.6$ | 7.638% |

**Figure 1:**
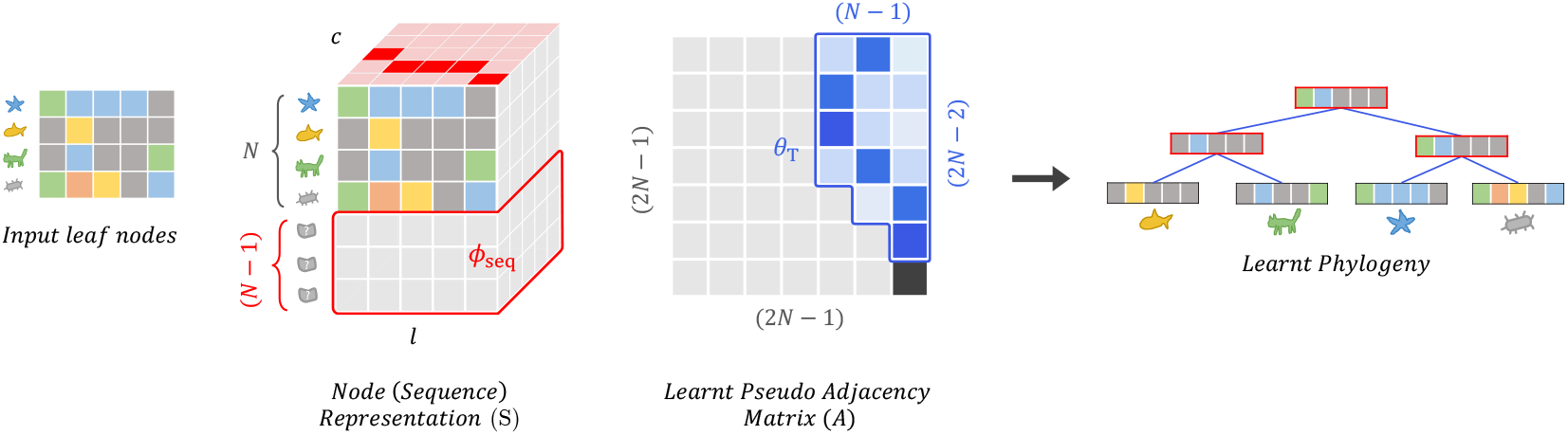
Visual depiction of the constructed adjacency matrix and the node representation using learnable parameters *θ*_*T*_ and *ϕ*_*seq*_.

**Figure 2:**
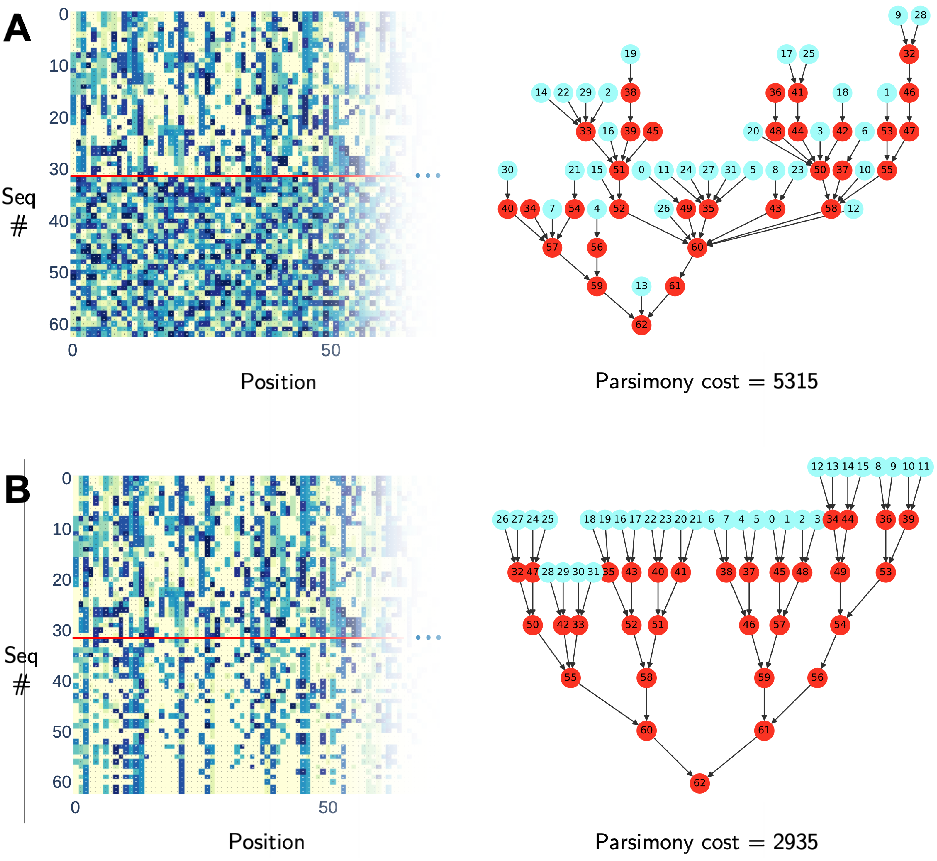
Search for evolutionary tree (*N* = 32, *l* = 256) left : sequences (leaves and ancestors), right : tree topology. A) random initialization, B) converged solution. optimal solution cost = 2913.

**Figure 3:**
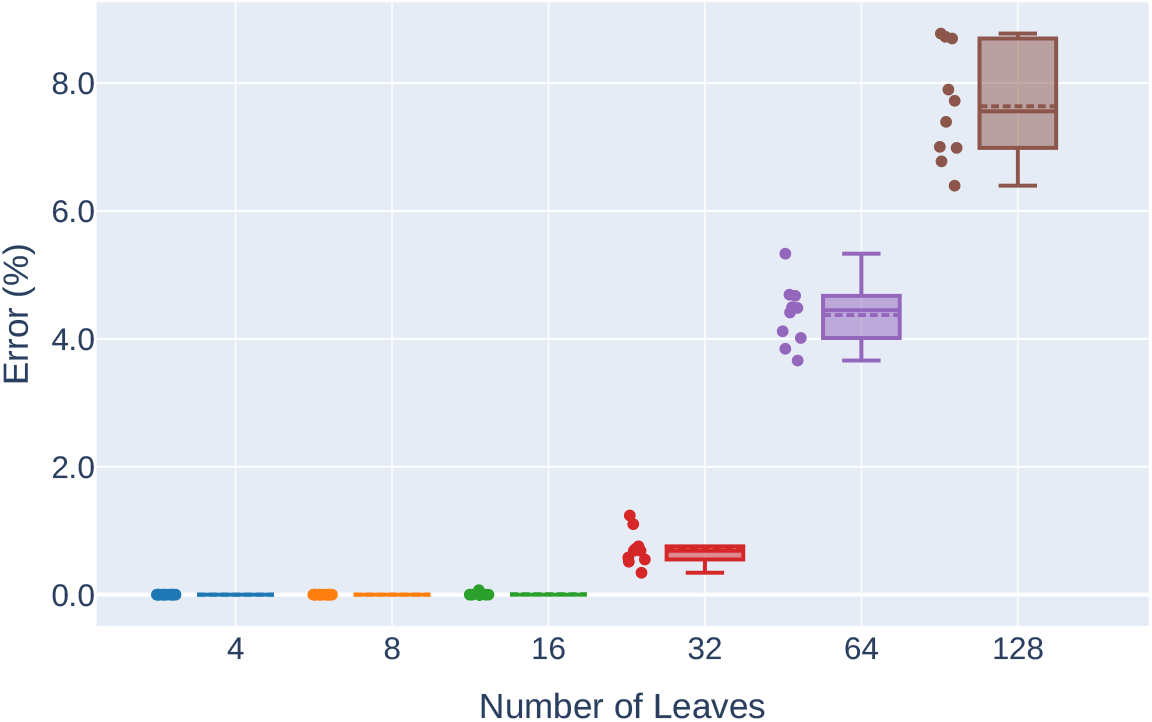
Evaluation on the maximum parsimony problem (find both the tree and ancestors, given leaves). Error is calculated w.r.t Sankoff algorithm solutions on the groundtruth tree topology.

**Figure 4:**
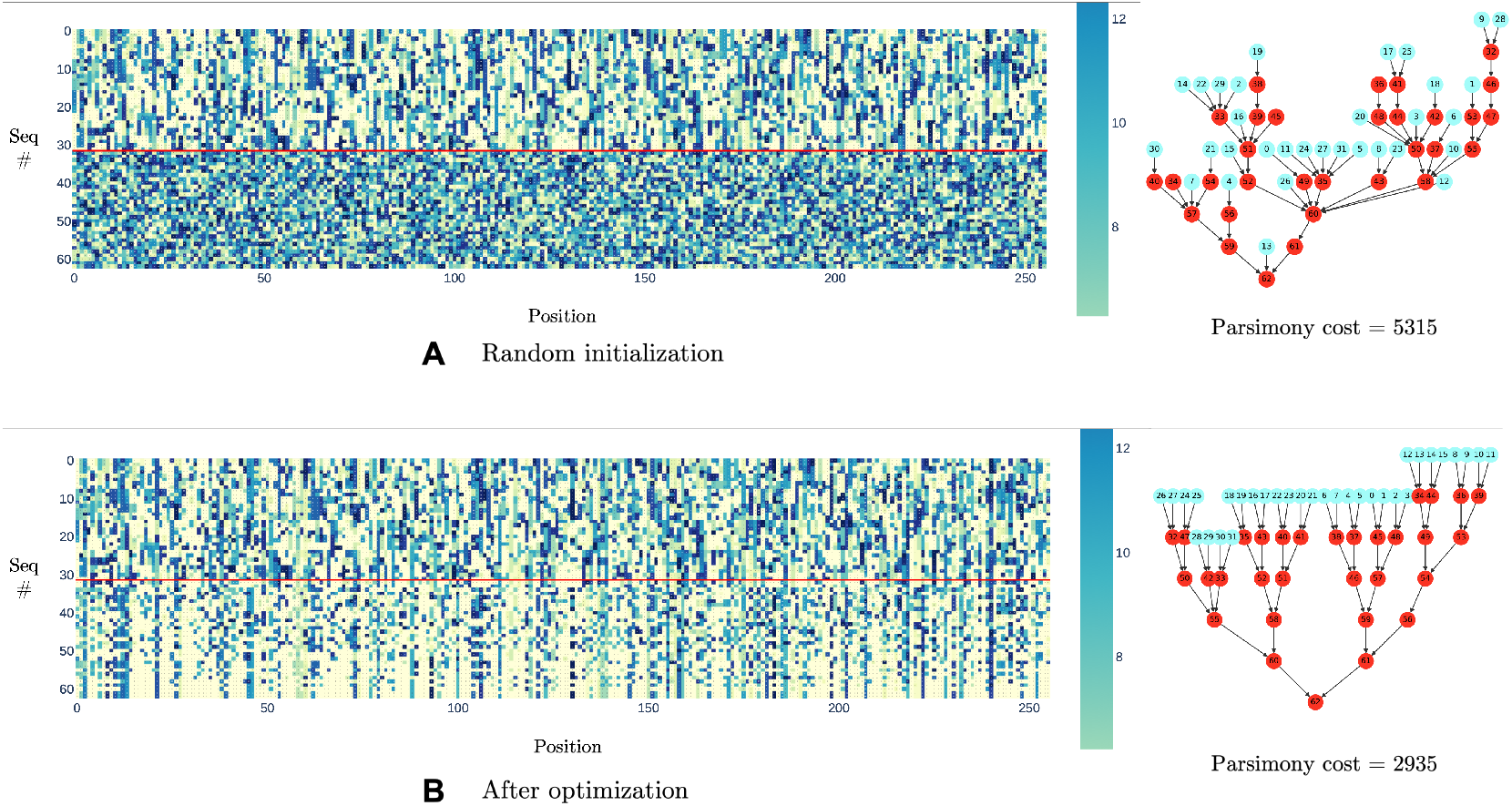
Search for evolutionary tree (*N* = 32, *l* = 256) left : sequences, right : tree topology. A) random initialization, B) converged solution. optimal solution cost = 2913.

### 4.3 Ablation of optimization methods

As mentioned in Section 3.4, we compare the bi-level optimization procedure with an independent alternative optimization scheme. In this formulation, we treat the tree parameters and sequence parameters independently and alternatively perform gradient descent using the Adam optimizer. We find that, even though this procedure works similarly well for trees with fewer leaves (*N ≤*16), it accumulates high error as the tree grows. The results are shown in Table 3.

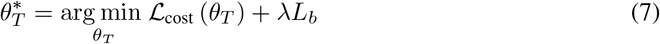

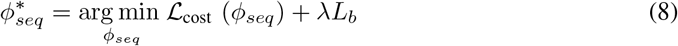

**Table 3:** Ablation study on two different optimization approaches on the maximum parsimony problem.

|  | Bi-level optimization |  |  | Alternative Optimization |  |  |
| --- | --- | --- | --- | --- | --- | --- |
| $N$ | Mean solution | Mean error | Mean error as a % w.r.t optimal solution | Mean solution | Mean error | Mean error as a % w.r.t optimal sol. |
| 4 | 277.2 | $0.0 \pm 0.0$ | 0.000% | 277.2 | $0.0 \pm 0.0$ | 0.000% |
| 8 | 653.1 | $0.0 \pm 0.0$ | 0.000% | 653.1 | $0.0 \pm 0.0$ | 0.000% |
| 16 | 1407.7 | $0.1 \pm 0.3$ | 0.007% | 1411.1 | $3.5 \pm 5.6$ | 0.249% |
| 32 | 2936.3 | $20.9 \pm 7.4$ | 0.717% | 2977.9 | $62.5 \pm 18.1$ | 2.144% |
| 64 | 6188.6 | $259.3 \pm 27.4$ | 4.373% | 6341.0 | $411.7 \pm 64.3$ | 6.943% |
| 128 | 12885.5 | $914.4 \pm 99.6$ | 7.638% | 13597.29 | $1632.0 \pm 129.0$ | 13.639% |

## 5 Discussion and Future Work

Our work establishes a new direction for generating evolutionary trees by traversing a soft tree and sequence space. Although here we focused on minimizing the parsimony cost as the objective (with unit cost for any change), our general optimization method can be coupled with various loss functions. For example, the parsimony cost assumes site-independence, which means that any position-wise evolutionary dependence in amino acids is ignored. Thus, if the tree is known, this independence property can be utilized to develop a dynamic programming algorithm that can derive the most optimal ancestral sequences (i.e. Sankoff algorithm Sankoff (1975)). Therefore, our method can be most beneficial when this condition is lifted (e.g. integration of pseudo-likelihood between nodes using protein language models or those conditioned on protein structure).

Even though the error increases as the number of leaves grows, future work could explore an iterative procedure to combine subtrees gradually (e.g. for 128 leaves, breaking down the problem to two sets of 64 leaves or 4 sets of 32 leaves and running gradient descent, then using these answers as a better initialization for the original problem).

Although our current work is primarily focused on solving the maximum parsimony problem (i.e. finding the parsimonious tree given only leaves), in the future, we plan to explore the application of our method within the context of Bayesian phylogenetic inference, inspired by recent advances in the field Swanepoel et al. (2022); Fourment et al. (2023); Zhang (2020); Zhang & Matsen (2019); Moretti et al. (2021).

## A Appendix

### A.1 Groundtruth data generation

We generate complete binary trees starting with a random sequence as root, make two copies of the sequence at each node, and generate two random sets of indices each with *m* = 50. These two sets of random indices are mutated into two copies, so that a new random amino acid is introduced at each index. It should be noted that this process does not necessarily mean that the generated complete binary trees are the trees with minimum number of evolutionary steps for the reached leaves (for these leaves there could be a better tree topology and ancestors). However, since mutations are introduced only at 50/256 ≈20% of the sequence length, the probability of the existence of better topologies is low, yet we find the best ancestors for this topology by applying the Sankoff algorithm, and this serves us as the groundtruth. Therefore, these serve as test samples to assess whether the optimization procedure converges to tree and its corresponding ancestors with evolutionary steps that are sufficiently close to the generated groundtruth.

**Figure 5:**
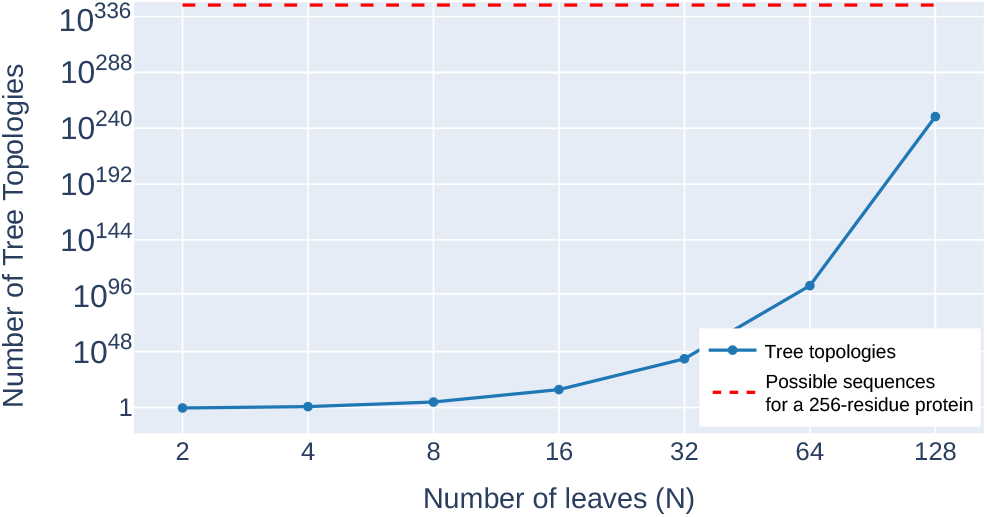
Number of rooted bifurcating tree topologies for *N* leaves (blue) and number of possible amino acid sequences (red) for a single 256-residue protein (i.e. 20^256^ ≈ 1.16 × 10^333^)

**Figure 6:**
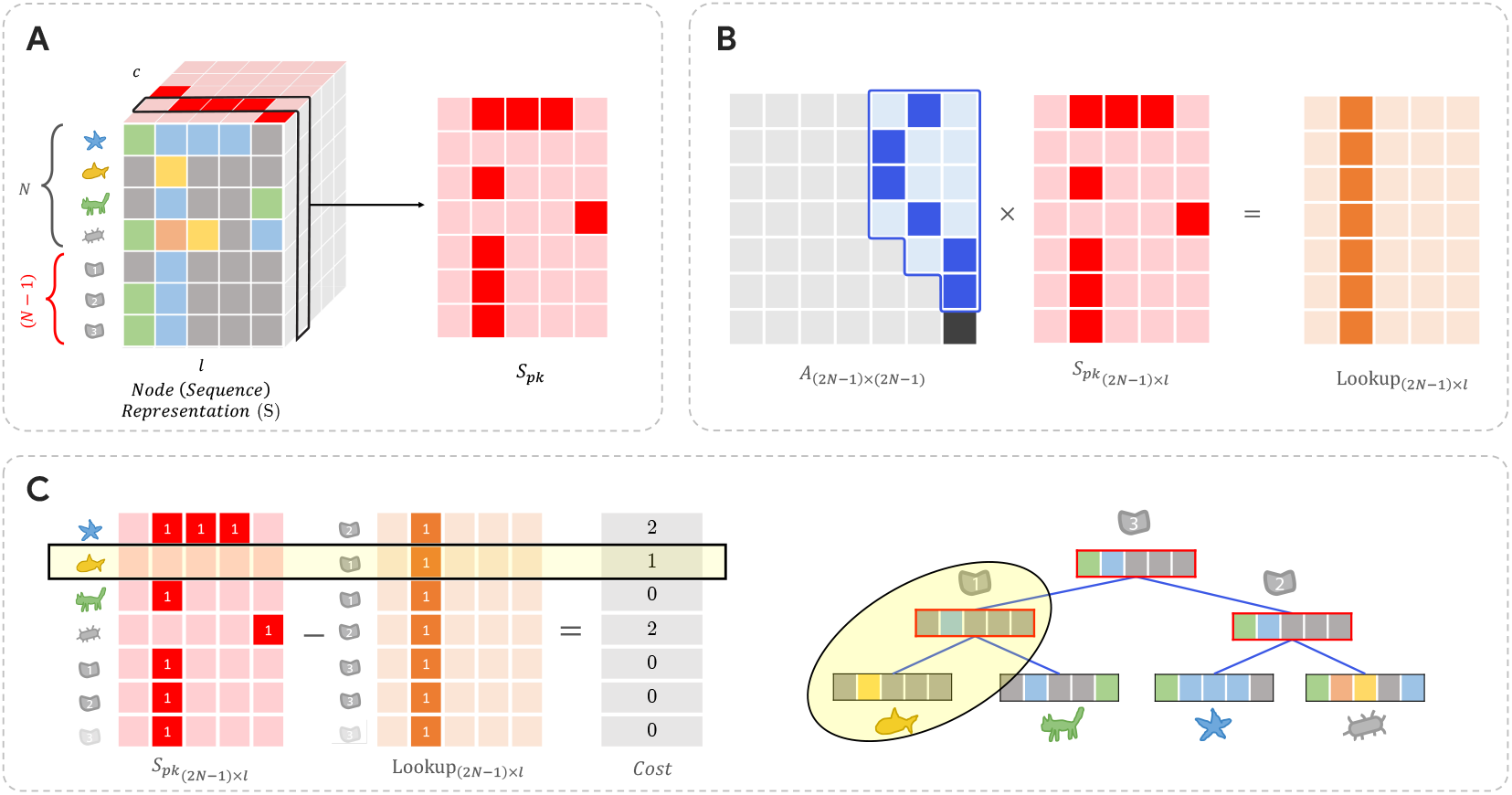
Explanation of parsimony cost calculation. A) considering one character dimension at a time. B) The matrix multiplication *A* × *S*_*pk*_ serves as a look-up of the sequence table (*S*_*pk*_). C) Difference between the child and parent sequence in the *k*^*th*^ character space.

## Notes

https://github.com/ramithuh/diff-evol-tree-search

### Competing Interest Statement

The authors have declared no competing interest.

